# Annual dynamics and distribution of *Xylella fastidiosa* in infected almond trees

**DOI:** 10.1101/2023.07.17.549336

**Authors:** Noa Zecharia, Vanunu Miri, Orit Dror, Kamel Hatib, Doron Holland, Shtienberg Dani, Ofir Bahar

**Affiliations:** Department of Plant Pathology and Weed Research, Agricultural Research Organization, Volcani Center, Rishon LeZion, Israel; The Robert H. Smith Faculty of Agriculture, Food and Environment, The Hebrew University of Jerusalem, Rehovot, Israel; Newe Ya’ar Research Center, Agricultural Research Organization, Volcani Center, Ramat Yishay, Israel

## Abstract

This research focused on studying the dynamics of the bacterial pathogen *Xylella fastidiosa* in almond trees at different developmental stages and in various tree parts. The objective was to understand the annual distribution and concentration of *X. fastidiosa* within almond trees. Different tree parts, including leaf petioles, annual and perennial shoots, fruit parts, flowers, and roots, from ten *X. fastidiosa*-infected almond trees were sampled over two years. The distribution and concentration of *X. fastidiosa* were determined using qPCR and serial dilution plating. Throughout the study, *X. fastidiosa* was never found in the fruit, flowers, and roots of almond trees, but it was present in leaves and annual and perennial shoots. We show that the inability of *X. fastidiosa* to colonize roots is likely due to incompatibility with the GF677 rootstock. The presence of *X. fastidiosa* in shoots remained consistent throughout the year, while in leaf petioles it varied across developmental stages, with lower detection during early and late stages of the season. Similarly, viable *X. fastidiosa* cells could be isolated from shoots at all developmental stages, while in leaf petioles no successful isolations were achieved during the vegetative and nut growth stage. Examining the development of almond leaf scorch symptoms over time in trees with preliminary infections revealed that once symptoms have appeared on a single branch, other asymptomatic limbs were likely already colonized by the bacterium, hence, selective pruning of symptomatic branches is unlikely to cure the tree. Overall, this study enhances our understanding of *X. fastidiosa* dynamics in almonds and may have practical applications for its detection and control in almond orchards.

## INTRODUCTION

*Xylella fastidiosa* is a plant pathogenic bacterium that depends on xylem-sucking insects for transmission (EFSA 2018; Hill and Purcell 1997). Strains belonging to the species *X. fastidiosa* are genetically diverse and belong to six different subspecies, which all together can colonize over 500 plant species (Delbianco et al. 2021). *X. fastidiosa* subspecies are commonly grouped into ‘sequence types’ (STs), which are determined by the nucleotide sequence of seven housekeeping genes (Scally et al. 2005). Strains belonging to a particular subspecies, or ST, can colonize and infect only a subset of the complete host range, however, the genetic factors determining the host specificity of *X. fastidiosa* remain mostly elusive.

Plant infection occurs when insect vectors that carry *X. fastidiosa* feed on leaf xylem, thereby introducing the bacterium into the vascular tissue of the plant. The spatial and temporal dispersion of *X. fastidiosa* from the infection site depends on the interactions between the specific pathogen strain and the host plant, as well as on environmental conditions (Feil and Purcell 2001). Disease symptoms generally appear weeks to months after infection and depend on the ability of the bacterium to proliferate and disperse in the host plant (White et al. 2020).

In compatible interactions, as with *X. fastidiosa* Temecula1 strain and grapevine, for example, pathogen cells move basipetally and acropetally within the xylem (Meng et al. 2005; Hill and Purcell 1995; de La Fuente et al. 2007), colonizing new shoots as well as perennial tissues (i.e., cordon and trunk). Substantial infection of the plant leads to xylem vessel clogging as a result of biofilm build up and the plant immune reaction to restrict pathogen spread (Sun et al. 2013; Newman et al. 2003). Clogging of the vessels disrupts the normal flow of water and often leads to leaf scorching, branch desiccation and die back symptoms, all of which are characteristic of *X. fastidiosa* diseases. These symptoms are most prominent during summer months when water demand by the plant is apparently not met by the limited water supplied through the occluded vessels (McElrone et al. 2001; Zecharia et al. 2022; Purcell et al. 1999).

*X. fastidiosa* infection of perennial plants can become chronic and persist for years. Persistence of the bacterium in perennial plants depends on the host species, cultivar (Purcell 1980b; Cao et al. 2011; Ledbetter et al. 2009), *X. fastidiosa* strain (Almeida and Purcell 2003), time of infection (Feil et al. 2003; Cao et al. 2011) and winter temperatures (Purcell 1980a). With respect to the time of inoculation, in both almonds and grapevines, plants that were infected early in the season had a higher likelihood of developing chronic infection. This is because the pathogen had sufficient time to reproduce and spread to the perennial tissues, allowing it to survive through the winter (Cao et al. 2011; Feil et al. 2003). On the other hand, plants that were infected later in the season had a greater chance of recovering from the infection. This phenomenon, where previously infected and symptomatic plants become uninfected in the following season, is known as ‘winter curing’ or ‘cold therapy’.

In 2017, *X. fastidiosa* subsp. *fastidiosa* (ST1) was reported to have infected almond (*Prunus dulcis*) orchards in northern Israel, causing almond leaf scorch (ALS) disease (Zecharia et al. 2022). Almond orchards throughout Israel are planted with locally bred cultivars, predominantly Um ElFahem (UEF, ~70 % of the cultivated area) and its pollinizers (‘Kochav’, ‘Gilad’, ‘Shefa’, and ‘Kochba’), grafted primarily on GF677, a peach-almond hybrid rootstock. Almond is a deciduous tree that sheds all, or most of it leaves during the dormancy stage, which in Israel occurs during the main winter months (December-January). Following the dormancy stage, flower buds swell, trees begin to bloom (February), and new leaves and shoots emerge (early March). From March to July, almond kernels mature and grow to full size and hull splits open. Trees are harvested by mechanical trunk shakers during peak summer months (August-September). In the weeks preceding harvest, irrigation is paused to minimize trunk damage due to shaking. Irrigation is resumed after harvest and the trees undergo another stage of vegetative growth. In November, leaves begin to senesce and fall in preparation for dormancy.

In Israel, most of the infected almond trees exhibit foliar symptoms on 50 % of the canopy or more, suggesting that they have been infected for some time (Zecharia et al. 2022). It is currently not clear how fast the bacterium, or ALS symptoms, progress in almond trees from the initial infection under commercial settings. Despite being chronically infected, new leaves re-emerge every spring with no visible symptoms and only later during summer ALS symptoms begin to appear. This cycle of symptoms appearance and disappearance in almonds is apparently governed by the deciduous nature of the trees. However, the timing of bacterial colonization on new leaves and its availability for vector acquisition is currently unclear. Understanding the spatial and temporal dynamics of *X. fastidiosa* during this cycle is essential for comprehending disease etiology, epidemiology, and disease diagnostics. The aim of this study was to characterize the dynamics and distribution of *X. fastidiosa* in various tree organs during different developmental stages along the year.

## MATERIALS AND METHODS

### Orchard and tree selection

Trials were carried out in two commercial almond orchards in Hula Valley in northern Israel: Kefar-Blum (33.171321, 35.598106) and Sde-Nehemia (33.195637, 35.630268). Trees were 13-14 years old and were grown under similar soil and climatic conditions as well as management practices. Ten almond trees of the cultivar UEF, (grafted on GF677 rootstocks) displaying ALS symptoms on >50 % of the canopy were chosen and labelled in 2018. Each tree represented one biological replicate and used for sampling and analysis as described below. Two asymptomatic almond trees from a commercial orchard (3-4 years old) in Sde-Nehemia, where ALS had not been detected, were chosen as negative controls. From June 2019 onwards, over the course of two consecutive years, the following tree organs were sampled: leaf petioles, annual shoots (green shoots that emerged in the same season) perennial shoots (lignified shoots that are at least one season old), roots, flower buds, flowers, and fruits (Fig. 1). Each sample was collected from four quadrant of the canopy of each tree and placed in a plastic bags. The samples were transferred to the laboratory in a cooling box and were subsequently kept at a temperature of 4°C for a maximum of 24 hours before undergoing further processing. Tree samplings were performed during the following developmental stages: blooming, vegetative and nut growth, hull split, harvest, leaf fall, and dormancy, during the respective months of the year as described in Table 1. Perennial and annual shoots as well as roots were sampled at all stages, while flowers and flower buds, fruits and leaves were sampled when present in the respective developmental stage.

**Figure 1.**
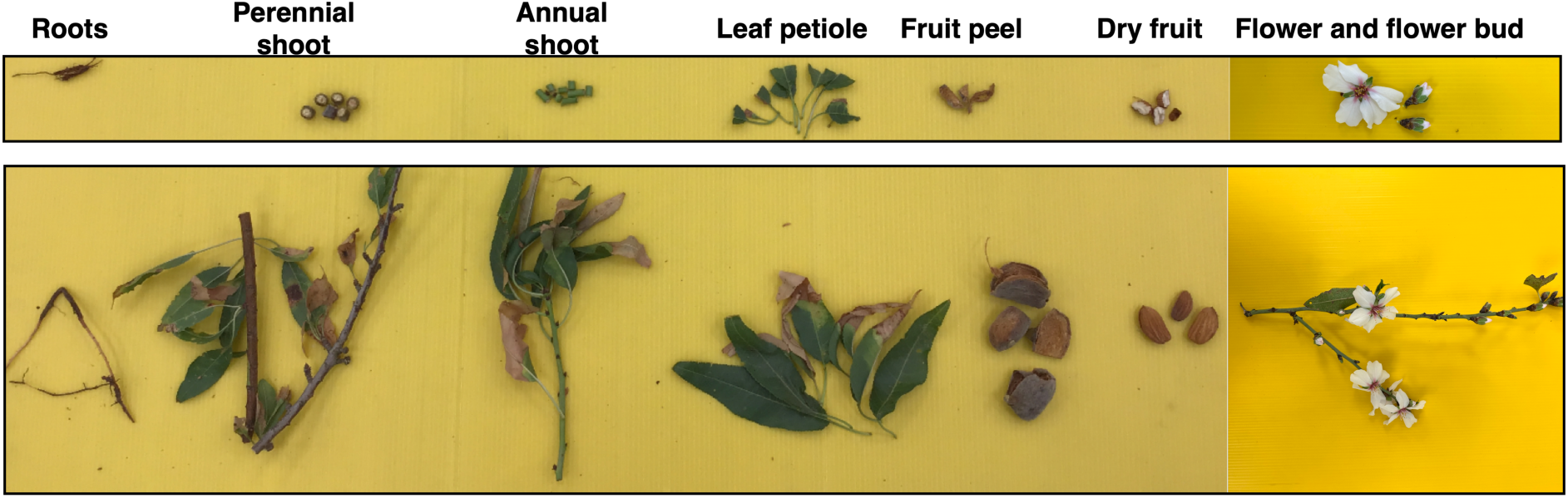
Almond organs and tissues collected for analyses. Roots, perennial and annual shoots, leaf petioles, fruit peel, dry fruit, flowers and flower buds were collected from almond trees at the relevant developmental stages and were used to determine *X. fastidiosa* incidence and relative titer by quantitative PCR, culturing and fluorescence *in situ* hybridization (FISH) analyses. Lower panel: samples as they were collected from the field and brought to the lab; Upper panel: processed tissue used for the different analyses.

**Table 1.** Almond developmental stages and their respective months of the year.

| Season | Developmental stage | Month of the year |
| --- | --- | --- |
| Winter | Blooming | February |
| Spring | Vegetative and Nut growth | March-May |
| summer | Hull split | June-July |
| Summer | Harvest | August |
| Fall | Leaf fall | November |
| Winter | Dormancy | December-January |

### DNA extraction from plant tissue

DNA was extracted from leaf petioles, annual and perennial shoots, flowers and flower buds using a modified CTAB (cetyltrimethylammonium bromide) procedure (Zhang et al., 1998), as previously described (Zecharia et al., 2021). In brief, 0.5 g of plant tissue were homogenized in Bioreba extraction bags using a semi-automated homogenizer (Homex 6, Bioreba, Switzerland) with 5 ml of CTAB buffer and 10 µl of 1M DTT (Dithiothreitol). Aliquots of the homogenized tissue (1.5 ml) were incubated at 65°C for 45 min with occasional mixing by vortex. The DNA extraction procedure continued as previously described (Mawassi et al. 2018).

DNA from root, fruit and fruit peel tissues was extracted using the DNeasy Mericon Food Kit (Qiagen). In brief, 0.5 g tissue was homogenized using the Homex 6 homogenizer as described above with 5 ml of Food lysis buffer (supplied), and DNA was subsequently purified according to the manufacturer’s instructions. DNA purity and concentration were determined for all samples using a Nanodrop device (Thermo Scientific, USA).

### Conventional and quantitative-PCR (qPCR)

Conventional PCR to identify *X. fastidiosa* was performed using the RST31/RST33 primer set (Minsavage et al. 1994) and 20 ng of template DNA. The DreamTaq Green PCR Master Mix (Thermo Scientific) was used in a final reaction volume of 10 µl with the following amplification conditions (denaturation: 95°C, 30 sec; annealing: 59°C, 30 sec; extension: 72°C, 1 min; X30 cycles). PCR products were analyzed on a 1 % agarose gel containing GelRed (0.01%, Biotium) and visualized under UV light.

Quantitative-PCR (qPCR) was performed with 20 ng of template DNA using the Fast SYBR Green Master Mix (Thermo Scientific) in a final reaction volume of 10 µl. The XF-F/XF-R primer set (Harper et al. 2010) was used to detect and quantify *X. fastidiosa*. For relative quantitation of *X. fastidiosa* titer, the almond actin gene was used as a normalizing gene (Forward primer PdActin-F: 5’-AGACCTTCAATGTGCCTGCT-3’; and Revers primer PdActin-R: 5’-AGCAAGGTCCAGACGAAGAA-3’). The relative *X. fastidiosa* concentration was calculated using the ΔCt method.

### Isolation and enumeration of *X. fastidiosa* cells from plant material

To isolate and enumerate *X. fastidiosa* from plant tissue, 0.5 g of leaves, annual or perennial shoots were washed thoroughly in soap and water and disinfected as previously described (Zecharia et al., 2021). The disinfected samples were homogenized in extraction bags as describe above with 5 ml PBS buffer (pH 7.4). Extract aliquots (1 ml) were transferred into a 1.7-ml Eppendorf tube and a series of ten-fold dilutions were made. Six 10-µl drops from each dilution were spotted on PD3 agar plates containing 40 mg/l cycloheximide and incubated at 28°C for 7-15 days until colonies suspected to belong to *X. fastidiosa* appeared. The number of colony forming units (CFU) per g tissue/developmental stage was determined by counting and averaging the number of colonies in ten biological replicates. A subset of the colonies was subjected to conventional PCR as described above to validate their identification.

### Rootstock and root colonization by *X. fastidiosa* in artificially inoculated almond plantlets

Plantlets were grown in 8-l pots in a net house and inoculated with *X. fastidiosa* (strains SN17-1, SN17-2, SN17-4, HY19-1) at 10^8^ CFU/mL using the needle prick method during April-May as described (Zecharia et al., 2021). In the following year, plantlets that tested positive for *X. fastidiosa* (by qPCR) were used for sampling of rootstock and roots (n=10), and healthy uninoculated plantlets were used as controls (n=3). The rootstock stem was sampled at two locations (25 cm below the grafting point and 25 cm above the soil surface) and used for DNA purification as described above. Roots were collected from the soil media and used for DNA purification using the DNeasy Mericon Food Kit, as described above. Finally, all samples were tested for *X. fastidiosa* by qPCR, as described above.

To determine whether *X. fastidiosa* can colonize and induce disease symptoms in the GF677 genotype, un-grafted GF677 plantlets were used (n=10). UEF plantlets (n=10) grafted on GF677 were used as controls. Plantlets were grown as described above and inoculated using the needle-prick method in five shoots per plantlet. Negative control plants of each genotype (n=3) were inoculated similarly with a 10-µl drop of sterile PD3 medium. Symptom development was monitored monthly for seven months post inoculation. *X. fastidiosa* incidence and titer were determined at five time points from June to November as described herein. At each time point, a single inoculated shoot was collected from each plantlet (n=10) and 0.5 g of leaf petioles were collected from the inoculation point (IP) and five-to-ten cm above it. Samples were immediately used for both qPCR analysis and CFU/g tissue determination, as described above.

### Fluorescence in situ Hybridizations (FISH) for *X. fastidiosa* detection in plant tissue

Infected and non-infected almond trees from commercial orchards (described above) were analyzed by FISH. From each tree (n=10), one annual and one perennial shoot with ALS symptoms were collected. One-mm-wide cross sections (n=5) were cut from leaf petioles, annual and perennial shoots using a sterile razor blade. Sections were placed in an Eppendorf tube and incubated overnight in fixation solution (Kliot et al. 2014). The next day, the fixation solution was removed, and sections were suspended for six hours in absolute ethanol at room temperature. Sections were washed three times with 1 ml of hybridization buffer (20 mM Tris-HCl, pH 8.0, 0.9 M NaCl, 0.01% sodium dodecyl sulfate, 30% formamide) before adding fresh hybridization buffer containing a final concentration of 1 ng/µl of fluorescently labeled probe KO 210 Cy3 (Cardinale et al. 2018). Sections were incubated in hybridization buffer with probe overnight at room temperature. The following day, sections were washed three times with fresh hybridization buffer, then placed on glass slides, and observed under an Olympus confocal laser scanning microscope (Tokyo, Japan) using wavelengths of 543 and 570 nm for excitation and emission, respectively.

### ALS symptom progress and *X. fastidiosa* distribution in almond trees with localized infection

In 2021-2022 we identified eight almond trees with preliminary ALS symptoms appearing on a single branch of the tree. The symptomatic branches were labelled, and the trees were monitored for ALS symptoms in the following year. Additionally, six of these trees were tested for *X. fastidiosa* by qPCR at: 1) symptomatic branches; 2) asymptomatic branches; and 3) base of each limb. Symptomatic and asymptomatic branches were tested by collecting leaf petioles from each branch and testing them by qPCR. The limb base was tested by drilling a hole (drill size = 10 mm) at the base of the limb (close to the joining point with the trunk) and collecting sawdust. DNA was extracted from sawdust using the CTAB procedure as described above. Trees were tested again in a similar manner in the following year but could not be followed further since the orchard was uprooted in winter 2022.

### Statistical analyses

The effect of ‘year of sampling’ and the effect of ‘year of sampling’ x ‘developmental stage’ were not statistically significant (*p* = 0.9957 and *p* = 0.3006, respectively); therefore, data from both years were pooled for analyses. The incidence of *X. fastidiosa* in different tree organs during different developmental stages was analyzed using the Pearson χ^2^ test. *X. fastidiosa* relative concentration as obtained by either qPCR (ΔCt method), or CFU counts, were analyzed by one-way analysis of variance (ANOVA), followed by the Tukey Kramer-HSD test, where applicable. Only samples positive for *X. fastidiosa* (Ct<35) were included in the ΔCt analysis.

## RESULTS

### Determining *X. fastidiosa* incidence and relative titer in different tree organs during different developmental stages using qPCR

Monitoring *X. fastidiosa* incidence in different organs of infected almond trees during different developmental stages was conducted during two consecutive years (2019-2021) by qPCR. Through all samplings, *X. fastidiosa* was never detected in roots, flowers, flower buds, fruits, or fruit peel of the infected trees. However, *X. fastidiosa* was detected in leaf petioles, and in annual and perennial shoots. Leaf petioles were collected for sampling during different developmental stages, excluding ‘dormancy’ and ‘blooming’ when no leaves were available. In leaf petioles, the incidence of *X. fastidiosa* was significantly lower in the ‘vegetative and nut growth’ (March-April) and ‘leaf fall’ (November-December) stages, compared with the ‘hull split’ and ‘harvest’ (June-August) stages (χ^2^= 44.304; df = 4; *p* < 0.001) (Fig. 2 and Table 1).

**Figure 2.**
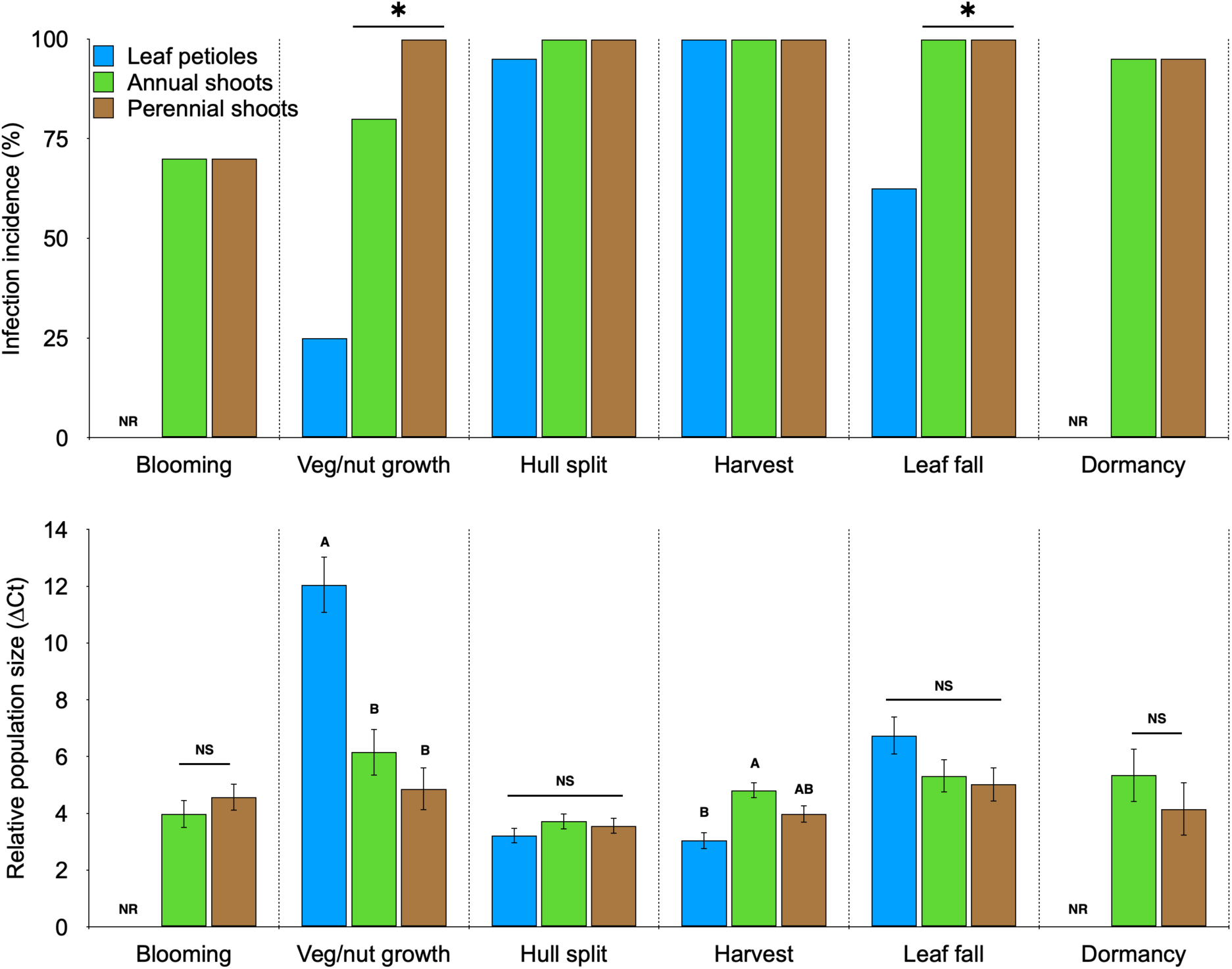
Determining *X. fastidiosa* incidence and relative titer in different tree organs during different developmental stages using qPCR. *X. fastidiosa* incidence (upper panel) and relative titer (lower panel) were tested in leaf petioles, annual and perennial shoots collected from infected almond trees in the orchard by qPCR during different developmental stages for two consecutive years. In the upper panel, asterisks indicate significant difference (*p* < 0.01) between *X. fastidiosa* incidence in leaf petioles versus annual and perennial shoots by the χ^2^ test (n=10). In the lower panel, *X. fastidiosa* relative titer was estimated by qPCR using the almond actin gene as a normalizer and calculating the ΔCt. Higher values indicate lower bacterial titer. Different letters indicate significant (*p* < 0.01) differences among tree organs by the Tukey-Kramer HSD test. NS, not significant; NR, not relevant (leaf petioles were absent during dormancy and blooming).

Annual and perennial shoots were sampled during all developmental stages and *X. fastidiosa* incidence in these tissues was more consistent than in leaf petioles. In perennial shoots, the incidence of *X. fastidiosa* among different developmental stages was not significantly different (p>0.05) excluding the ‘blooming’ stage (χ^2^ = 26.245; df = 5; *p* < 0.0001). The mean incidence of *X. fastidiosa* in perennial shoots throughout all sampling timepoints was 94.17 ± 4.56 % (mean ± SD). In annual shoots, *X. fastidiosa* incidence was significantly lower during ‘blooming’ and ‘vegetative and nut growth’ stages compared with the other developmental stages (χ^2^ = 19.716; df = 5; *p* < 0.0014) (Fig. 2). The mean incidence of *X. fastidiosa* throughout all sampling timepoints was 90.83 ± 4.95 %. When comparing the incidence of *X. fastidiosa* among the different organs during different developmental stages we found that the incidence of *X. fastidiosa* in annual and perennial shoots was significantly higher than in leaf petioles in the ‘vegetative and nut growth’ (χ^2^ = 27.882; df = 2; *p* = 0.0001) and the ‘leaf fall’ (χ^2^ = 10.909; df = 2; *p* = 0.0043) stages (Fig. 2).

The relative titer of *X. fastidiosa* in leaf petioles during different developmental stages varied significantly (one-way ANOVA; *F*_3,70_ = 32.1739, *p* = 0.001). A post hoc Tukey-Kramer HSD test indicates that the relative titer of *X. fastidiosa* during the ‘hull split’ and ‘harvest’ stages was significantly (*p* < 0.01) higher than in the other developmental stages. *X. fastidiosa* titer in petioles was the lowest in the ‘vegetative and nut growth’ stage (*p* = 0.0001) (Fig. 2). When comparing the relative titer of *X. fastidiosa* in annual and perennial shoots in different developmental stages, we found no statistical differences (one-way ANOVA; *F*_5,104_ = 1.7817, *p* = 0.1230 and *F*_5,107_ = 0.8393, *p* = 0.5247, for annual and perennial shoots, respectively). When comparing the relative titer of *X. fastidiosa* among different tree organs, significant differences were found in the ‘vegetative and nut growth’ (one-way ANOVA; *F*_2,60_ = 17.3228, *p* = 0.0001) and ‘harvest’ (one-way ANOVA; *F*_2,48_ = 9.3273, *p* = 0.0004) stages. A post hoc Tukey-Kramer HSD test indicates that the relative titer of *X. fastidiosa* in the ‘vegetative and nut growth’ stage, was significantly higher in both annual and perennial shoots compared with leaf petioles (*p* = 0.0001), however, it was significantly lower in annual shoots compared with leaf petioles during the ‘harvest’ stage (*p* = 0.0002) (Fig. 2).

### Isolation of *X. fastidiosa* cells from different tree organs during different developmental stages

The concentration of viable *X. fastidiosa* cells in petioles and stems of infected almond trees was estimated during different developmental stages by tissue maceration and serial dilution plating. *X. fastidiosa* cells could be isolated from leaf petioles, and from annual and perennial shoots, at all the developmental stages tested. Successful isolation from leaf petioles was significantly higher during ‘hull split’ (80 %) and ‘leaf fall’ (45 %) compared with the ‘vegetative and nut growth’ stage (χ^2^= 17.400; df = 2; *p* < 0.001), where isolation success rate was zero (Fig. 3). The average concentrations of *X. fastidiosa* in petioles in the ‘hull split’ and ‘leaf fall’ stages were 6.42 and 6.2 log10 CFU/gr, respectively (Fig. 3). The isolation success rate from annual and perennial shoots was not statistically different among developmental stages (annual shoots, χ^2^= 1.54; df = 3; *p* = 0.7641; perennial shoots, χ^2^ = 4.371; df = 3; *p* = 0.2241) and had an average of 55 ± 4 % and 38 ± 10 %, respectively. The average log10 values of *X. fastidiosa* CFU from annual and perennial shoots per gram tissue were 5.93 and 5.32, respectively (Fig. 3). Among the different organs, there was a significantly higher chance of recovering *X. fastidiosa* from annual shoots compared with leaf petioles in the ‘vegetative and nut growth’ stage (χ^2^ = 8.571; df = 2; *p* < 0.034). On the other hand, chances to recover *X. fastidiosa* from leaf petioles were significantly higher (χ^2^ = 10.179; df = 2; *p* < 0.001) than annual and perennial shoots during ‘hull split’ (summer), however, the concentration of recovered cells was not statistically significant among them (one-way ANOVA; *F*_2,29_ = 0.8506, *p* = 0.4376).

**Figure 3.**
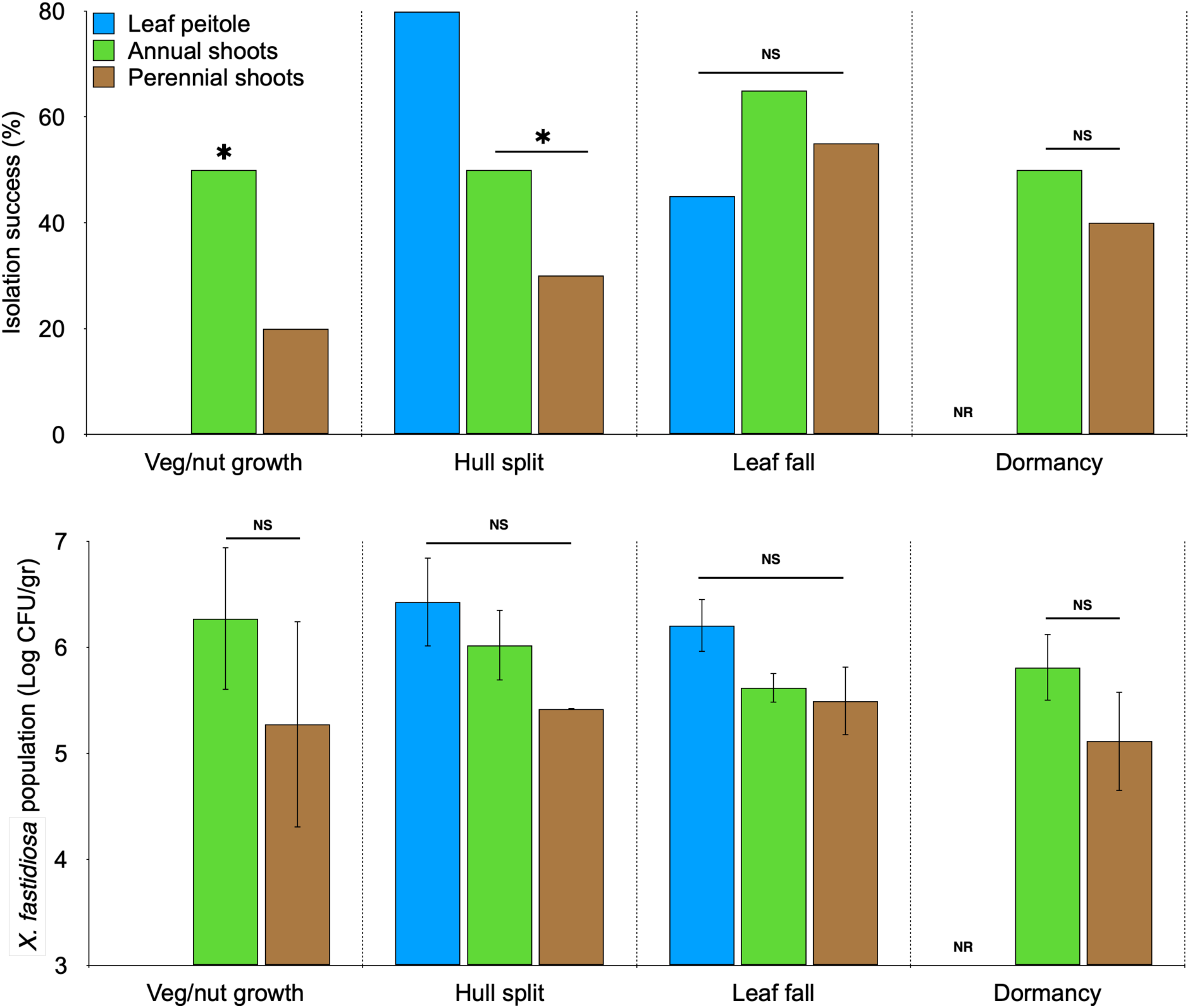
Isolation of *X. fastidiosa* cell from different tree organs during different developmental stages. *X. fastidiosa* isolation success (upper panel) and CFU per gram tissue (lower panel) from different tree organs through different developmental stages were determined. Tree organs were sampled as described in Fig. 1 (n=10). Asterisks in the upper panel indicate significant differences compared with leaf petiole at each developmental stage (χ2 test, *p* < 0.01). In the lower panel, *X. fastidiosa* CFU per gram in the different organs was statistically not significant (NS) in any of the developmental stages tested (ANOVA, *p* > 0.05).

### FISH analysis of *X. fastidiosa* in different tree organs during different developmental stages

Cross and longitudinal sections of petioles and shoots of infected and control almond trees were examined during the ‘hull split’ and ‘leaf fall’ stages under a confocal microscope using a *X. fastidiosa* specific probe. During ‘hull split’, when ALS symptoms are clearly visible, *X. fastidiosa* cells could be detected in 97.9 % of the petiole sections examined, while during ‘leaf fall’, the incidence of positive sections was 31.8 % (Table 2). Aggregates of *X. fastidiosa* were detected within xylem vessels in both cross and longitudinal sections of infected petioles, whereas no such aggregates were observed in petioles from control trees (Fig. 4). In annual shoots, *X*. *fastidiosa* detection rate with FISH was 10 to 12 % in ‘hull split’ and ‘leaf fall’ stages, whereas in perennial shoots, the bacterium could not be detected using FISH in any of tested samples (Table 2).

**Figure 4.**
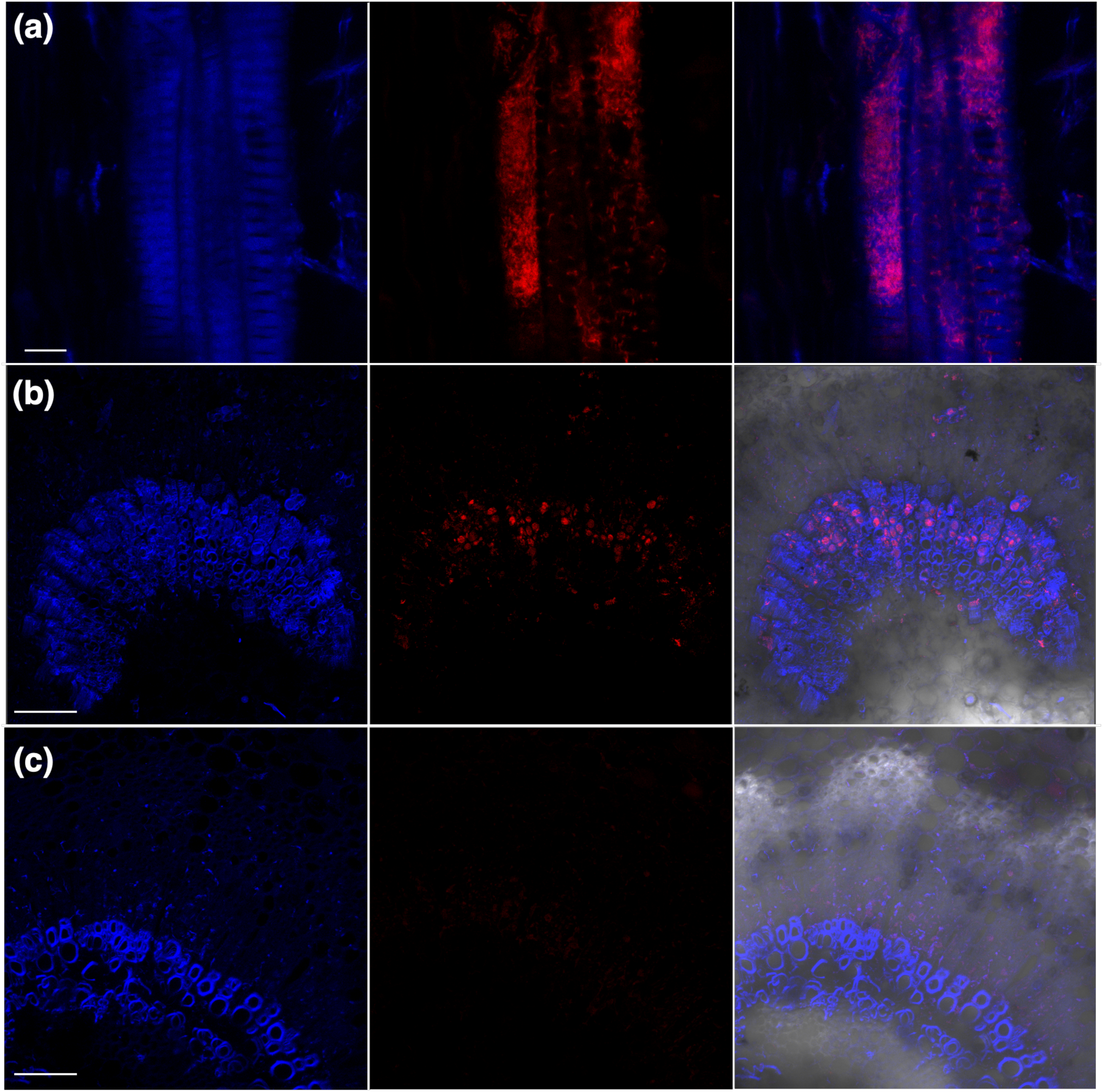
Detection of *X. fastidiosa* in almond leaf petioles using FISH. Longitudinal (**a**) and cross (**b**, **c**) sections of almond leaf petioles were incubated with a *X. fastidiosa*-specific Cy3 probe (red signal) and examined under a confocal microscope. Leaf petioles were collected from symptomatic (**a**, **b**) and asymptomatic control (**c**) trees in the orchard. Images in each panel are of the same specimen, showing plant autofluorescence (left, blue), *X. fastidiosa* Cy3 probe fluorescence (middle, red) and a merged image (right). Scale bar represents 10 µm for panel (a) and 100 µm for panels (b) and (c).

**Table 2.** Fluorescence *in situ* hybridization (FISH) of *X. fastidiosa* in different tree organs in two different developmental stages.

| Developmental stage | Hull split |  | Leaf fall |  |
| --- | --- | --- | --- | --- |
|  | No. of specimens tested (positive/total) | Positive specimens (%) | No. of specimens tested (positive/total) | Positive specimens (%) |
| Leaf petiole | 46/47 | 97.87 | 14/44 | 31.82 |
| Annual shoots | 6/50 | 12 | 4/49 | 10.2 |
| Perennial shoots | 0/14 | 0 | 0/36 | 0 |

### Rootstock and root colonization by *X. fastidiosa* in artificially inoculated almond plantlets

*X. fastidiosa* was not detected in either GF677 rootstock stems or roots of all ten infected plantlets (Table 3). *X. fastidiosa* was detected by qPCR at the inoculation point (IP) of GF677 and UEF plantlets at similar rates, however, it was detected five-to-ten cm above the IP primarily in the UEF genotype (Fig. 5). Furthermore, *X. fastidiosa* was successfully isolated from UEF plantlets from both the IP and five-to-ten cm above it at multiple time points post inoculation, while it could not be isolated from any of the GF677 inoculated plantlets (Fig. 5). Disease symptoms were not observed four months post inoculation in the GF677 inoculated plantlets, while 70 % of the inoculated UEF plantlets were symptomatic at the same time.

**Figure 5.**
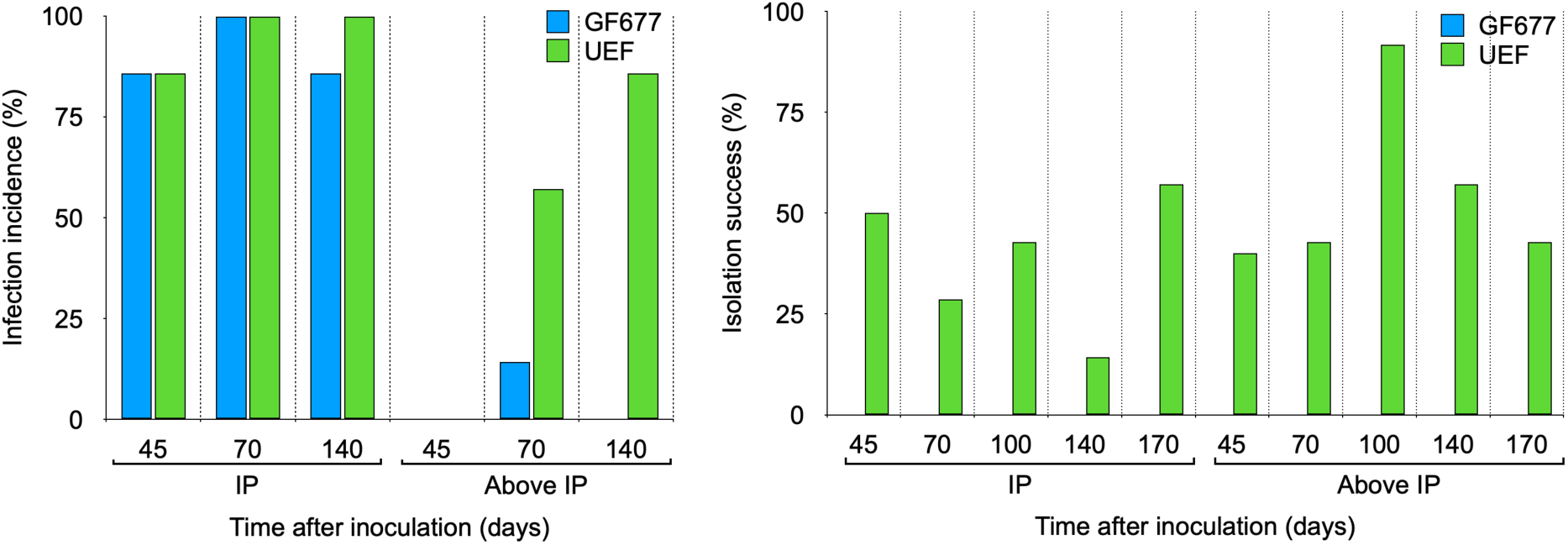
*X. fastidiosa* incidence by qPCR and isolation from artificially inoculated UEF and GF677 plantlets. UEF and GF677 plantlets were mechanically inoculated with *X. fastidiosa* and then sampled at different time points post inoculation from the inoculation point (IP) or above (5 to 10 cm). Left panel describes *X. fastidiosa* incidence at the IP or above, as determined by qPCR. Right panel describes *X. fastidiosa* isolation success by plating on PD3 agar plates.

**Table 3.** *X. fastidiosa* incidence in rootstock and roots of mechanically inculcated almond plantlets.

| Replicate | Plantlet variety | Inoculated/control | qPCR result (Ct)* |  |  |  |
| --- | --- | --- | --- | --- | --- | --- |
|  |  |  | Scion (UEF) stem | Rootstock GF677_I** | Rootstock GF677_II*** | GF677 Roots |
| 1 | UEF on GF677 | Inoculated | 30.05 | Neg | Neg | Neg |
| 2 | UEF on GF677 | Inoculated | 27.10 | Neg | Neg | Neg |
| 3 | UEF on GF677 | Inoculated | 31.88 | Neg | - | Neg |
| 4 | UEF on GF677 | Inoculated | 29.84 | Neg | - | Neg |
| 5 | UEF on GF677 | Inoculated | 29.14 | Neg | - | Neg |
| 6 | UEF on GF677 | Inoculated | 33.42 | Neg | Neg | Neg |
| 7 | UEF on GF677 | Inoculated | 31.33 | Neg | Neg | Neg |
| 8 | UEF on GF677 | Inoculated | 29.57 | Neg | - | Neg |
| 9 | UEF on GF677 | Inoculated | 29.31 | Neg | - | Neg |
| 10 | UEF on GF677 | Inoculated | 30.05 | Neg | Neg | Neg |
| 11 | UEF on GF677 | Control | Neg | Neg | Neg | Neg |
| 12 | UEF on GF677 | Control | Neg | Neg | Neg | Neg |
| 13 | UEF on GF677 | Control | Neg | Neg | Neg | Neg |

\*, Ct values above 35 were considered negative; \*\*, rootstock stem sampled 25 cm below the graft point; \*\*\*, rootstock stem sampled 25 cm above the soil surface (rootstock stem was ~ 1-m long)

### ALS symptom and *X. fastidiosa* progression in trees with localized infection

Eight almond trees with localized ALS symptoms (appearing only on a single tree limb) were identified and labelled in 2021. Leaves from symptomatic limbs were tested by qPCR and found positive for *X. fastidiosa*, while leaves from asymptomatic limbs were mostly negative (excluding one tree). In the following year, ALS symptoms spread to two or more additional limbs in 75 % of the trees (6/8, Table 4). When examining the sawdust extracted from limb base, 50 (3/6) and 17 % (3/17) of the symptomatic and asymptomatic limbs, respectively, were positive in 2022.

**Table 4.**
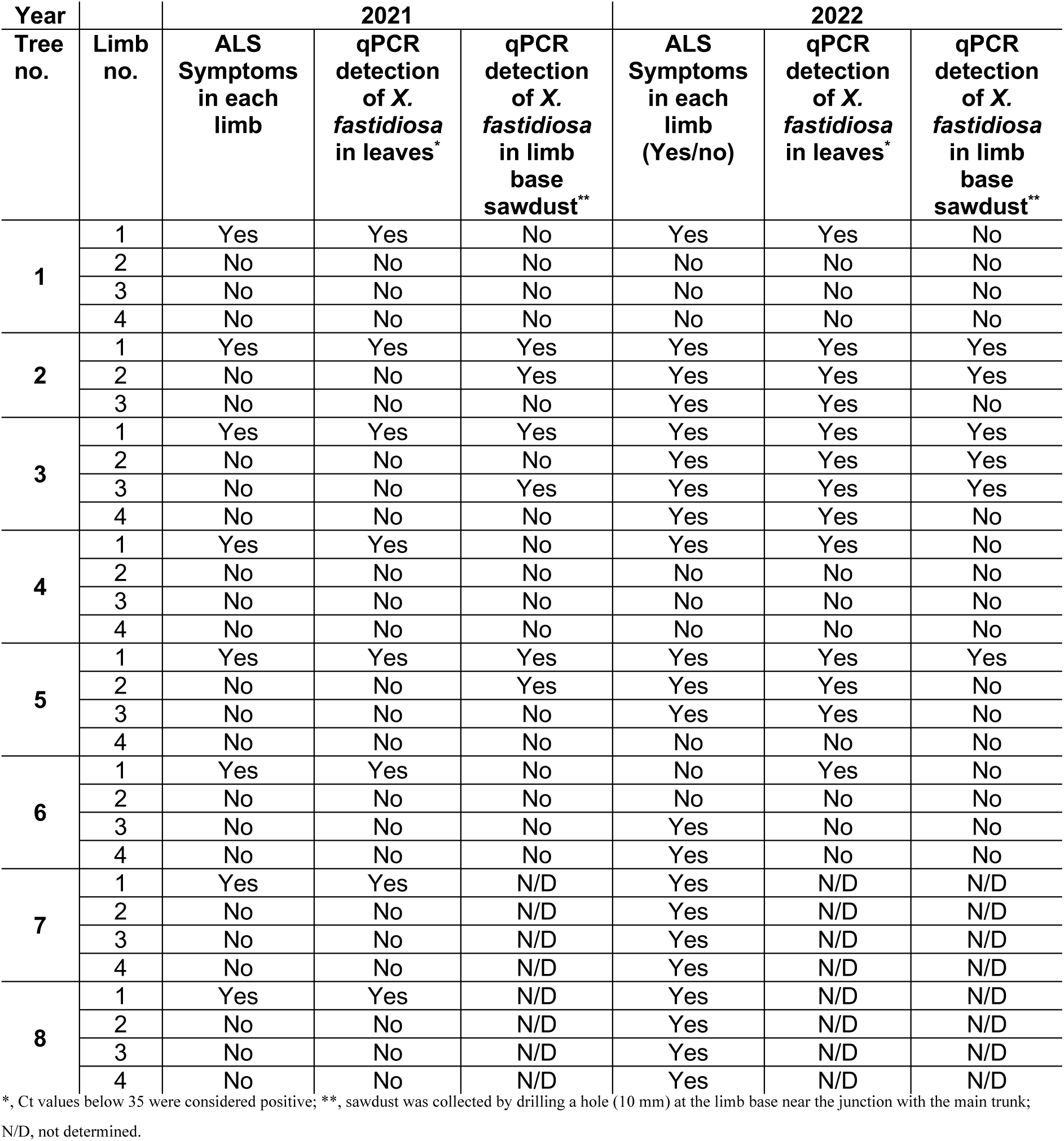
ALS symptoms and *X. fastidiosa* progression in trees with localized infection.

## DISCUSSION

While being chronically infected with *X. fastidiosa*, every spring almond trees develop new leaves that appear asymptomatic and healthy. The typical ALS symptoms only begin to appear during early summer months (June) and are most noticeable during August-September. This phenomenon differentiates between deciduous trees and evergreen perennials such as citrus and olive, where symptoms are visible throughout the whole year. We were therefore prompt to study how *X. fastidiosa* adapts to the deciduous cycle of almond trees, by following the annual dynamics of bacterium colonization and distribution in infected trees.

Upon leaf emergence during early spring, *X. fastidiosa* cells have to migrate from their overwintering sites, shoots and branches, into the emerging leaves. Our results indicate that in the ‘vegetative and nut growth’ stage, *X. fastidiosa* cannot be isolated from leaves of infected trees and remains mostly undetected by qPCR. The temperatures during this stage are relatively low and not conducive for bacterial replication and spread, which may explain the low incidence of *X. fastidiosa* in the leaves. A previous study has shown that a minimum of 10^4^ CFU/gr tissue is required for acquisition and transmission of *X. fastidiosa* by the blue-green sharpshooter (*Graphocephala atropunctata*) in grapevine (Hill and Purcell 1997). Together with our results, it indicates that *X. fastidiosa* is most likely beneath the threshold required for acquisition during this stage. During ‘hull split’, we saw a significant increase in the incidence and cell titer of *X. fastidiosa* in the leaves. This was reflected in the ability to successfully isolate viable *X. fastidiosa* cells in the range of 10^6^ CFU per gram of leaf petiole, suggesting that trees serve as a good inoculum source for insect acquisition at this stage. As trees enter dormancy, the occurrence and concentration of *X. fastidiosa* in leaves decrease. This reduction can be attributed to leaf senescence and bacterial death, as well as the migration of viable cells into perennial tissues where they survive throughout the winter (Rapicavoli et al. 2018). Consequently, these factors likely contribute to lower rates of pathogen acquisition during this stage. These results are in accordance with the report of Amanifar et al. (2016) from Iran, depicting a similar cycle of *X. fastidiosa* in almond leaves. Altogether, these results may have important implications for ALS disease epidemiology, as they suggest that *X. fastidiosa* acquisition and transmission by insect vectors, does not occur through the ‘vegetative and nut growth’ stage, but mainly during the ‘hull split’ and ‘harvest’ stages (June to September). These results also indicate that disease diagnostics in almonds based on leaves during the ‘vegetative and nut growth’ stage is unreliable and may be misleading.

Unlike in leaves, *X. fastidiosa* incidence in perennial tissues such as annual and perennial shoots was consistent through different developmental stages and on average was over 90 %. This indicates that *X. fastidiosa* can be molecularly detected in perennial tissues of almonds with high confidence throughout the whole year. This result has important implications for disease diagnostic efforts, which have been mostly focused on leaves during specific time of the year. Nevertheless, despite the ability to detect *X. fastidiosa* using qPCR in perennial tissues, the isolation rate of viable cells from these tissues was less consistent and generally lower than qPCR detection. It is not clear whether this is a mere technical difficulty to extract bacterial cells from hard and lignified tissue, or a result of a non-viable state of the bacteria in these organs. Our FISH analysis highlights the technical difficulties in visualizing *X. fastidiosa* cells in perennial tissues, as cells were visualized only in 12 % and 0 % of the specimens from annual and perennial shoots, respectively, in the ‘hull split’ stage. On the other hand, viable cells could be isolated from these tissues in the ‘hull split’ stage at much higher rates (30 to 50 %) indicating that the difficulty is more technical rather than a result of lack of viable cells during this developmental stage.

Interestingly, throughout this study we were unable to detect *X. fastidiosa* in tree organs such as flowers, fruits, and roots. *X. fastidiosa* was previously reported to translocate to and colonize the roots of perennial plants such as olive (dos Santos et al. 2022; Saponari et al. 2017), almond (Amanifar et al. 2016), Peach (Davis et al. 1981; Aldrich et al. 1992), blueberry (Holland et al. 2014), Sycamore (Henneberger et al. 2004) and citrus (Hopkins et al. 1991; He et al. 2000), and it was therefore somewhat surprising that we were unable to detect it in our study. Our previous study shows that *X. fastidiosa* strains from Israel do not infect peach, plum and other perennial crops (Zecharia et al. 2022). Since the almond trees tested in this study were grafted on GF677 rootstocks, which is a peach-almond hybrid, we suspected that the rootstock does not allow *X. fastidiosa* colonization and migration towards the roots. Direct inoculation experiments with GF677 plantlets revealed that *X. fastidiosa* cells could not effectively migrate away from the inoculation point in GF677 plantlets, unlike in UEF grafted on GF677. Additionally, while the majority of UEF plantlets displayed disease symptoms, GF677 plantlets did not. These results are in line with a previous study where artificial inoculations of peach-almond hybrids with *X. fastidiosa* (strain M23) did not lead to ALS symptoms, nor was the pathogen detected by qPCR (Ledbetter and Rogers 2009). Overall, this suggests that GF677 inherited genetic elements conferring *X. fastidiosa* immunity from peach as other peach-almond hybrids, rendering them non-hosts for *X. fastidiosa*. Nevertheless, while resistant rootstocks occasionally do confer enhanced resistance to the scion, in this particular case it appears that chronic and severe infection of almonds, as seen in almond orchards in Israel, can occur in cultivars grafted on non-host rootstocks despite the inability of the pathogen to colonize the root system. Further support to this claim comes from citrus, where it was shown that bacterial translocation from the aerial parts to the root system occurs but is not essential to induce *X. fastidiosa* foliar symptoms, and that citrus variegated chlorosis (CVC) leaf symptom severity is not correlated with bacterial translocation to the roots (He et al. 2000).

Symptoms progression under natural conditions is difficult to assess. Since most of the infected trees were already exhibiting ALS symptoms on large parts of the canopy, it was difficult to determine how fast the bacterium and ALS spreads since the initial infection. Luckily, we have identified a small number of trees in which symptoms appear only on a single limb. Our results indicate that in these trees the bacterium has not yet established in the leaves of the asymptomatic limbs, however, it is possible that it has already migrated to the base of the symptomatic limb and even to the base of asymptomatic limbs. Together with the fact that initial infections are very difficult to detect, these results imply that removal of symptomatic almond branches infected with *X. fastidiosa* is not likely to be a practical methodology to remediate tree infection, as was also seen with PD in grapevines and with CVC (Daugherty et al. 2018; Coletta-Filho et al. 2000). Furthermore, our results show that in trees with preliminary ALS symptoms (appearing on a single branch), in most cases, the symptoms will spread to additional branches and limbs of the tree within one season. While this does not answer the question of how long it takes for symptoms to develop since the initial infection under natural conditions, it does reveal that under the tested conditions, once ALS symptoms have appeared, the progression of symptoms in the canopy is expected to be rapid.

In summary, our results provide insights to the temporal and spatial dynamics of *X. fastidiosa* in almond trees, which follows this model: as almond trees exit dormancy, overwintering bacterial cells begin to migrate from the perennial tissues towards the developing leaves. Significant leaf colonization by *X. fastidiosa* is achieved approximately two months after leaf emergence and the pathogen titer continues to increase with time, peaking during mid-to late summer, when foliar symptoms are most clearly visible. As trees approach dormancy and leaves senesce, *X. fastidiosa* titer in the leaves decreases, probably due to bacterial death and migration to perennial tissues, where pathogen cells will hibernate until the next spring. These results together with vector acquisition and transmission experiments, may allow to define a critical window in time at which *X. fastidiosa* acquisition and transmission would most likely occur in almonds. The knowledge gained from these insights could assist in determining the optimal timing for implementing vector control measures in the orchard, thereby potentially reducing the spread of the disease.

## ACKNOWLEDGEMENTS

The authors wish to thank the almond growers, Mr. Yaron Pinto and Mr. Nir Glosman, for their collaboration during these experiments, and to Mr. Menahem Borenstein for his valuable advice and technical help. This research was funded by grant no. 20-02-0115 from the Chief Scientist of the Ministry of Agriculture and Rural Development of Israel and the Fruit Council.

